# Herpesviral induction of germline transcription factor DUX4 is critical for viral gene expression

**DOI:** 10.1101/2021.03.24.436599

**Authors:** Stephanie Walter, Vedran Franke, Nir Drayman, Emanuel Wyler, Savaş Tay, Markus Landthaler, Altuna Akalin, Armin Ensser, Florian Full

**Affiliations:** Institute for Clinical and Molecular Virology, University Hospital Erlangen, 91054 Erlangen, Germany; Berlin Institute for Medical Systems Biology, Max Delbrück Center for Molecular Medicine, Helmholtz Society, Berlin, Germany; The Pritzker School for Molecular Engineering, The University of Chicago, Chicago, IL, USA, 60637; Institute of Virology, Medical Center - University of Freiburg, Freiburg, Germany

## Abstract

DUX4 is a transcription factor and a master regulator of embryonic genome activation (EGA). During early embryogenesis, EGA is crucial for maternal to zygotic transition at the 8-cell stage in order to overcome silencing of genes and enable transcription from the zygotic genome. In adult somatic cells, DUX4 expression is largely silenced. Activation is likely pathogenic, and in adult muscle cells causes genetic disorder Facioscapulohumeral Muscular Dystrophy (FSHD).

We identified activation of DUX4 expression upon lytic replication of the herpesviruses HSV-1, HCMV, EBV and KSHV, but not of adenoviruses, negative strand RNA viruses or positive strand RNA viruses. We demonstrate by RNA-Seq analysis that DUX4 expression upon herpesviral replication leads to the induction of hundreds of DUX4 target genes including germline-specific retroelements as well as several members of the TRIM, PRAMEF and ZSCAN protein families. Moreover, we show that DUX4 expression is a direct consequence of herpesviral infection. DUX4 can be stimulated by overexpression of HSV-1 immediate early proteins, indicating active induction of EGA genes by herpesviral infection. We further show that DUX4 expression is critical for driving HSV-1 gene expression.

Our results show that viruses from alpha-, beta- and gamma-herpesvirus subfamilies induce DUX4 expression and downstream germline-specific genes and retroelements. We hypothesize that herpesviruses induce DUX4 expression in order to induce an early embryonic-like transcriptional program that prevents epigenetic silencing of the viral genome and facilitates herpesviral gene expression.

## Introduction

Herpesviruses are a major health burden worldwide, with a prevalence of 60 – 100% dependent on the virus species and geography^1–3^. Herpesviruses cause a number of prevalent diseases, like oral and genital herpes, chickenpox, shingles and infectious mononucleosis. While they rarely induce life-threatening infections in healthy humans, infections in e.g. immunocompromised patients, transplant recipients or newborns can be severe^4,5^. Some herpesviruses such as Epstein-Barr virus (EBV) and Kaposi’s sarcoma-associated herpesvirus (KSHV) are also known to be the causative agent of particular cancer types^6^. Due to their lifelong persistence in the host, the risk of reactivation and developing an lytic infection is constantly present. Treatment options are limited and therapies able to counteract the viral persistence or to clear the virus out of the body are unavailable. A better understanding of the mechanisms involved during herpesviral infection is essential.

In a recent study, we observed that the upregulation of the cellular protein Tripartite-motif 43 (TRIM43) upon HSV-1 infection is dependent on the embryonic transcription factor DUX4^7^, which was recently confirmed by Friedel et al. ^8^. DUX4 is a germline transcription factor exclusively expressed from the 4-cell to the 8-cell state, a short phase during human embryonic development^9–11^. There, DUX4 is essential for embryonic genome activation (EGA, equivalent to zygotic genome activation in other mammals)^10–12^, a transcriptional activation event that leads to the induction of hundreds of target genes and allows the embryo to proceed further in development^13,14^. Concomitantly, it induces retroelements such as LTRs and ERVs^9,10,15^ which function as alternative promotors, regulate EGA-related genes and mediate pluripotency^16–18^. After this limited period of activation, DUX4 is silenced throughout all adult tissues, except for spermatocytes^19^. An exception to this is the genetic disorder Facioscapulohumeral Muscular Dystrophy (FSHD). In FSHD, DUX4 silencing is lost and aberrant DUX4 expression in adult muscle cells leads to apoptosis and degeneration of the affected muscles^20–23^. The reasons for the onset of this disease are complex and not yet fully understood, though they are thought to lie within the unique genomic location of DUX4^22,24^. It resides within the macrosatellite repeat array D4Z4^25,26^, that harbors up to 100 copies of DUX4^27^, with only the most distal copy encoding for a protein^21^. During early embryonic development, this macrosatellite array is transformed into heterochromatin after the 8-cell stage^9,10^ and its repression is strictly controlled by chromatin remodeling factors and repressors which have not been fully elucidated ^19,24,28^.

Given the peculiar expression pattern of DUX4 and its upregulation in herpesviral infection, we aimed to determine whether DUX4 plays a role during herpesviral infection and to shed light on its potential functions. Our data reveal aspects of a mechanism in herpesviral infection that has hitherto not been characterized. We show that DUX4 expression during herpesviral replication is a general mechanism that is conserved among all herpesviral subfamilies. For HSV-1, we could demonstrate that the viral proteins ICP0 and ICP4 promote DUX4 expression and hundreds of DUX4 target genes, including several classes of endogenous retroelements, and that DUX4 is critical for HSV-1 gene expression. This points at so far unknown endogenous functions of DUX4 and extends our knowledge of this unique transcription factor.

## Results

We previously showed that DUX4 protein is expressed upon lytic infection with HSV-1^7^. To confirm the expression of DUX4 and DUX4 target genes TRIM49 and ZSCAN4 upon HSV-1 infection, we probed their expression by Western blot analysis in infected primary human foreskin fibroblast (HFF) cells and 293T cells (Fig. 1A-C). Next, we investigated whether this DUX4 expression is unique for HSV-1 or a general feature of herpesviral infection. We therefore assessed the expression of DUX4 upon infection with one member of the α-, β–, and γ-herpesvirus subfamilies, namely HSV-1, HCMV and KSHV respectively. Upon HCMV infection of primary HFF cells, DUX4 as well as ZSCAN4 protein expression could be robustly detected by Western Blot (Figure 1D and E). In addition, DUX4 mRNA was strongly increased upon infection with HSV-1 and HCMV, as well as lytic reactivation of KSHV (Fig. 1 F-H). Moreover, we could also detect induction of known DUX4 target genes TRIM48, TRIM49, ZSCAN4, ZSCAN5a, ZSCAN5d and RFPL4A ^29,30^ for all three viruses (Figure 1I-K, Fig. S1). This indicates that DUX4 is functional and acts as a transcriptional activator in herpesvirus infected cells. DUX4 expression was also induced upon infection of primary human melanocytes with HSV-1, emphasizing the physiological relevance of DUX4 induction (Fig. S2). In addition, reanalysis of several published next generation sequencing datasets from HSV-1 infected cells confirmed that observation^7,31,32^. Comparison of three independent RNA-seq datasets in 3 different cell types showed no expression of DUX4 and DUX4 target genes (e.g. FRG1) in uninfected cells (Fig 2A). Upon infection with HSV-1, however, transcription is induced for DUX4 as well as all known DUX4 target genes (Fig. 2A). Interestingly, we also confirmed the expression of HSAT-II satellite repeat transcripts. HSAT-II repeats are known DUX4 targets^12,33^ and have recently been shown to play a role in HCMV replication^34^. (Fig. S3) Moreover, the reanalysis of a published Precision Run-On sequencing (PRO-seq.) dataset^34^ showed no RNA-polymerase-II occupancy at the DUX4 locus in uninfected cells, indicating that the locus is completely silenced under normal conditions (Fig. 2A). In contrast, HSV-1 infection induced a robust PRO-seq signal at the DUX4 locus. Taken together, these data show that the induction of DUX4 expression after herpesviral infection is a common feature for human herpesviruses and is conserved among α-,β-, and γ-herpesviruses.

**Figure1:**
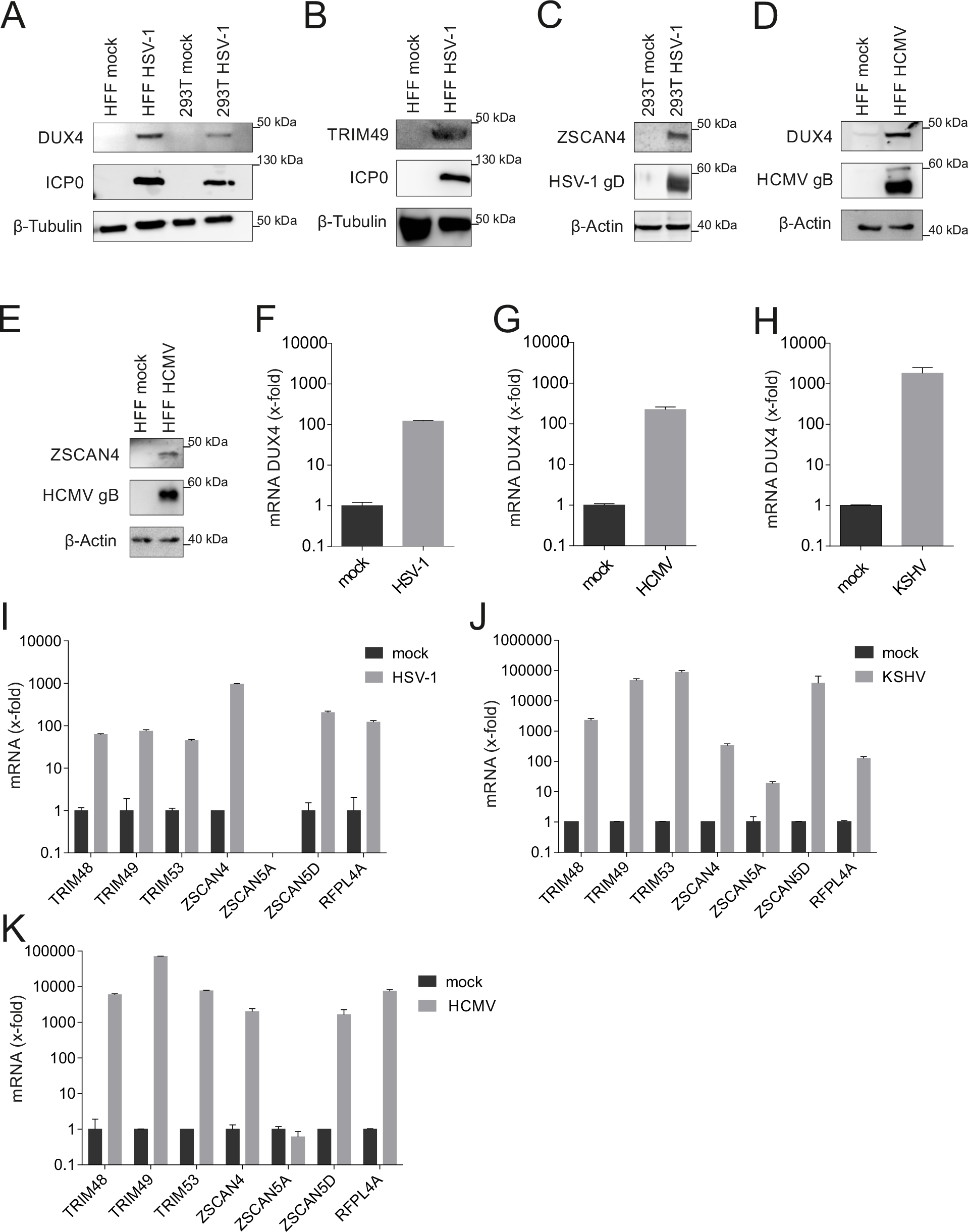
Expression of DUX4 and DUX4 target genes upon infection with α-, β-, γ-herpesviruses **A.** Western blot of primary HFF cells infected with HSV-1 for 24 h and 293T cells infected with HSV-1 for 18 h, both MOI 5. ICP0 was used as marker for infection. **B.** Western blot of HFF cells infected with HSV-1 for 24 h with MOI 5. **C.** Western blot of 293T cells infected with HSV-1 for 18 h with MOI 5. **D,E.** Primary HFF cells infected with HCMV for 6d with MOI 1. Western blot analysis of DUX4 (D) and ZSCAN4 (E). HCMV gB was used as marker for infection. **F-H.** qPCR analysis of DUX4 expression upon infection of 293T cells with HSV-1 MOI 5 for 18 h **(F)**, upon expression of HFF cells with HCMV MOI 1 for 6d **(G)** and upon lytic reaction of KSHV in iSLK cells at 5d p.i. **(H) I.** mRNA expression of DUX4 target genes in 293T upon infection with HSV-1 for 18 h with MOI 10. **J.** mRNA expression of DUX4 target genes in HFF infected with HCMV for 6 d with MOI 1. **K.** mRNA expression of DUX4 target genes in iSLK cells with reactivated KSHV infection, harvested after 5 d. In all qPCR experiments mRNA of targets was normalized to HPRT1 mRNA expression, one representative out of at least n=3.

**Figure 2:**
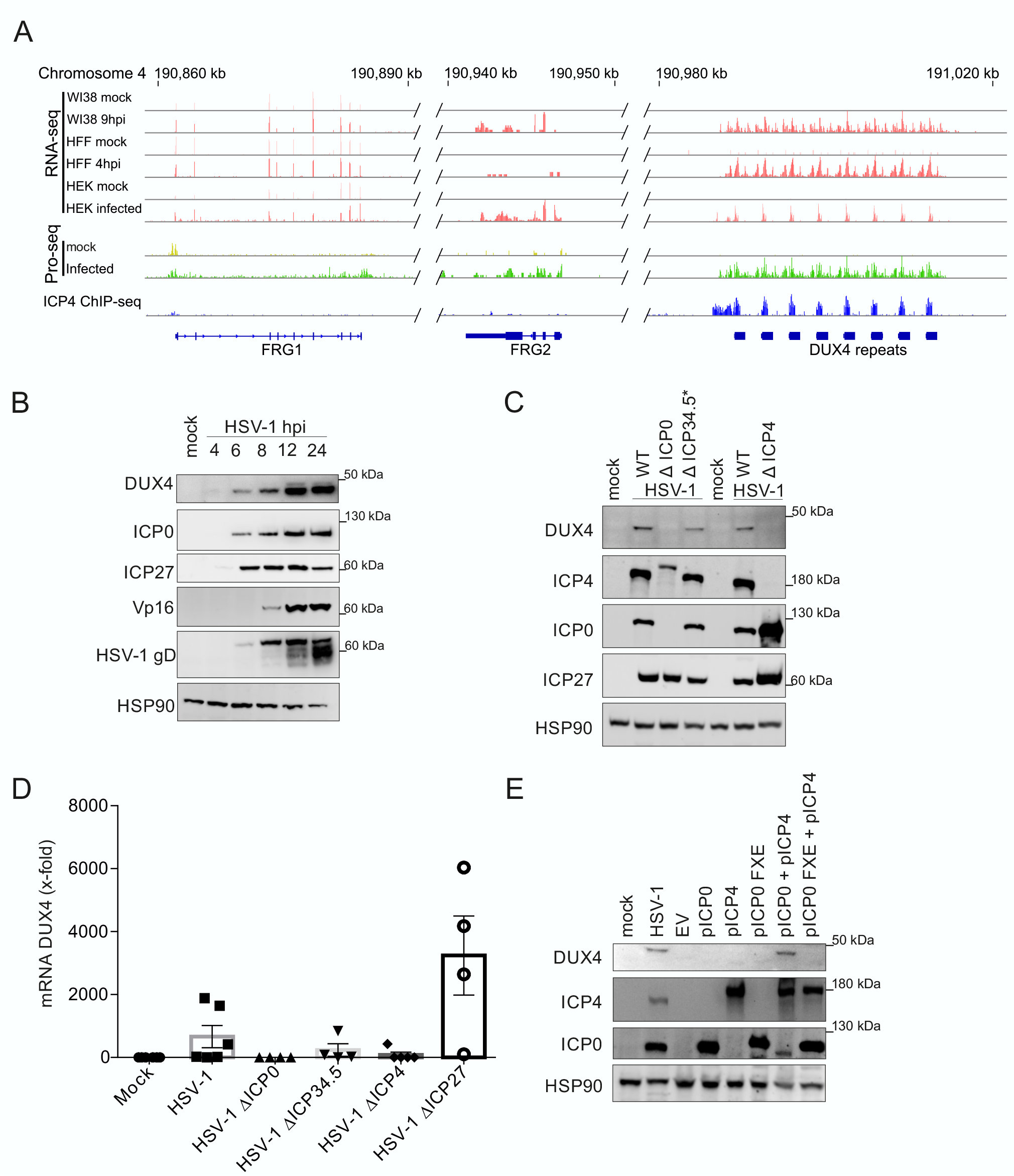
**A.** RNA-Seq. data of the DUX4 and neighbouring FRG1 and FRG2 loci in WI38 9 hpi, HFF 4 hpi and HEK 293T cells 18 hpi. The Precision Run-On Sequencing (Pro-Seq.) data show RNA POL II occupancy on the FRG1, FFRG2 and DUX4 locus in infected and uninfected cells. ICP4 ChIP-Seq. data show ICP4 occupation at the DUX4 locus in cells infected with HSV-1. **B.** Western Blot of DUX4 and HSV-1 proteins in primary HFF infected with HSV-1 harvested at different hpi (MOI 10). **C.** Western Blot of DUX4 and HSV-1 proteins in primary HFF infected with HSV-1 and HSV-1 mutants harvested at 16 hpi (MOI 10). One representative experiment out of n=5. **D.** qRT-PCR of DUX4 expression relative to HPRT from primary HFF cells infected with HSV-1, HSV-1 ΔICP0 (ICP4-YFP), HSV-1 ΔICP34.5, HSV-1 ΔICP4 for 24 h (MOI 10). **E.** Western blot analysis of DUX4 and HSV-1 protein in 293T cells transfected with HSV-1 IE proteins ICP0, ICP0 FXE, ICP4 (ICP4-YFP), ICP0 + ICP4 (ICP4-YFP) and ICP0 FXE + ICP4 (ICP4-YFP) for 48 h or infected with HSV-1 for 18 h (MOI 10). ICP0 FXE is a mutant with a deletion in the RING domain, which inhibits Ubiquitin E3 ligase activity. EV: empty vector control. One representative experiment out of n=3.

Next, we wanted to address the significance of herpesviral DUX4 induction for the infection itself, since DUX4 is a germline transcription factor that is not expressed in healthy adult tissue^9,10,19^. We first hypothesized that the induction of DUX4 might be part of an antiviral response of the cell to viral infection triggered by a herpesviral pathogen associated molecular pattern (PAMP), or by herpesviral induction of a cellular DNA-damage response (DDR). We reanalyzed several published RNA-sequencing datasets, investigating DUX4 mRNA expression in response to cellular changes, but found no evidence for DUX4 expression. In addition, triggering a DNA-damage response by treating cells with Bleocin or Etoposide did not induce DUX4 expression (Fig. S4A). The only evidence for DUX4 expression came from a Chromatin-IP-sequencing (ChIP-Seq.) dataset^36^ investigating the binding of the HSV-1 infected cell protein 4 (ICP4) to the cellular genome. ICP4 is a potent activator of viral transcription, part of the viral tegument, and essential for replication of HSV-1^37,38^. Our reanalysis showed a very strong binding of the ICP4 protein to the entire DUX4 locus (Figure 2A). Of note, the peak intensity of ICP4 binding at the DUX4 locus was among the highest in the entire host genome. This ICP4 binding was a strong indication for an active induction of DUX4 by HSV-1 infection. Moreover, experiments with phosphonoacetic acid (PAA), an inhibitor of the viral DNA-polymerase, showed that herpesviral DNA replication is dispensable for DUX4 expression (Fig. S4B), indicating that induction of the DUX4 gene takes place at the immediate-early stage of the viral gene expression cascade. Furthermore, after infection with UV-inactivated virus, DUX4 protein was still induced, indicating that incoming components of the virion contribute to DUX4 induction (Fig. S4B). Analysis of the kinetics of DUX4 expression upon HSV-1 infection further demonstrated that DUX4 protein could be detected as early as 4h post infection with protein levels constantly increasing over the course of infection (Fig. 2B). Reanalysis of RNA-sequencing data from Rutkowski et al. further demonstrated that known DUX4 target genes have variable dynamics during infection with most DUX4 target genes being upregulated at 3-4 hours post infection (Fig. S5). To further elucidate the involvement of HSV-1 tegument proteins in the induction of DUX4 expression we performed infection experiments with HSV-1 mutants depleted of the immediate early proteins ICP0, ICP4, ICP27 and gamma-34.5. We could show that wildtype (wt) HSV-1 infection as well as infection with HSV-1-Δ γ34.5 resulted in DUX4 expression, whereas DUX4 expression is abrogated in cells infected with HSV-1 lacking either ICP0 or ICP4 (Fig. 2C and D). In addition, the transient co-expression of ICP0 and ICP4 is sufficient to induce DUX4 expression in the absence of viral infection (Fig. 2E), confirming that ICP0 and ICP4 are necessary and sufficient for inducing DUX4 expression.

DUX4 is a transcription factor active in the nucleus where it executes its physiological function by activating its target genes. We used a recombinant HSV-1 expressing ICP4-YFP and VP26-RFP in order to visualize progression of infection and localization of DUX4. ICP4 is an immediate-early gene and part of the tegument, whereas VP26, as the small capsid protein of the virus, is expressed late in the infection cycle. By co-staining of cells with a DUX4 antibody, we could demonstrate that only ICP4+/VP26-cells (YFP+-/RFP-cells) show nuclear expression of DUX4, indicating activation of DUX4 within early stages of infection (Figure 3a). In contrast, ICP4+ and VP26+ cells (YFP+ and RFP+ cells) show either no DUX4 expression or an aberrant cytoplasmic localization of DUX4. This indicates that DUX4 is only briefly activated by ICP0 and ICP4 during the early phase of infection. To test this, we treated cells with PAA, which arrests viral infection at the stage of viral DNA replication, i.e. after immediate-early gene expression but before late gene expression (Figure 3B). PAA treated cells showed a strong increase in expression of DUX4 and its target genes by qRT-PCR analysis of HDF cells infected with HSV-1 (Figure 3B), as well as the DUX4 protein by WB analysis in 293T cells (Figure S4B), supporting the notion of a transient activation of DUX4 during the early stages of HSV-1 infection.

**Figure 3:**
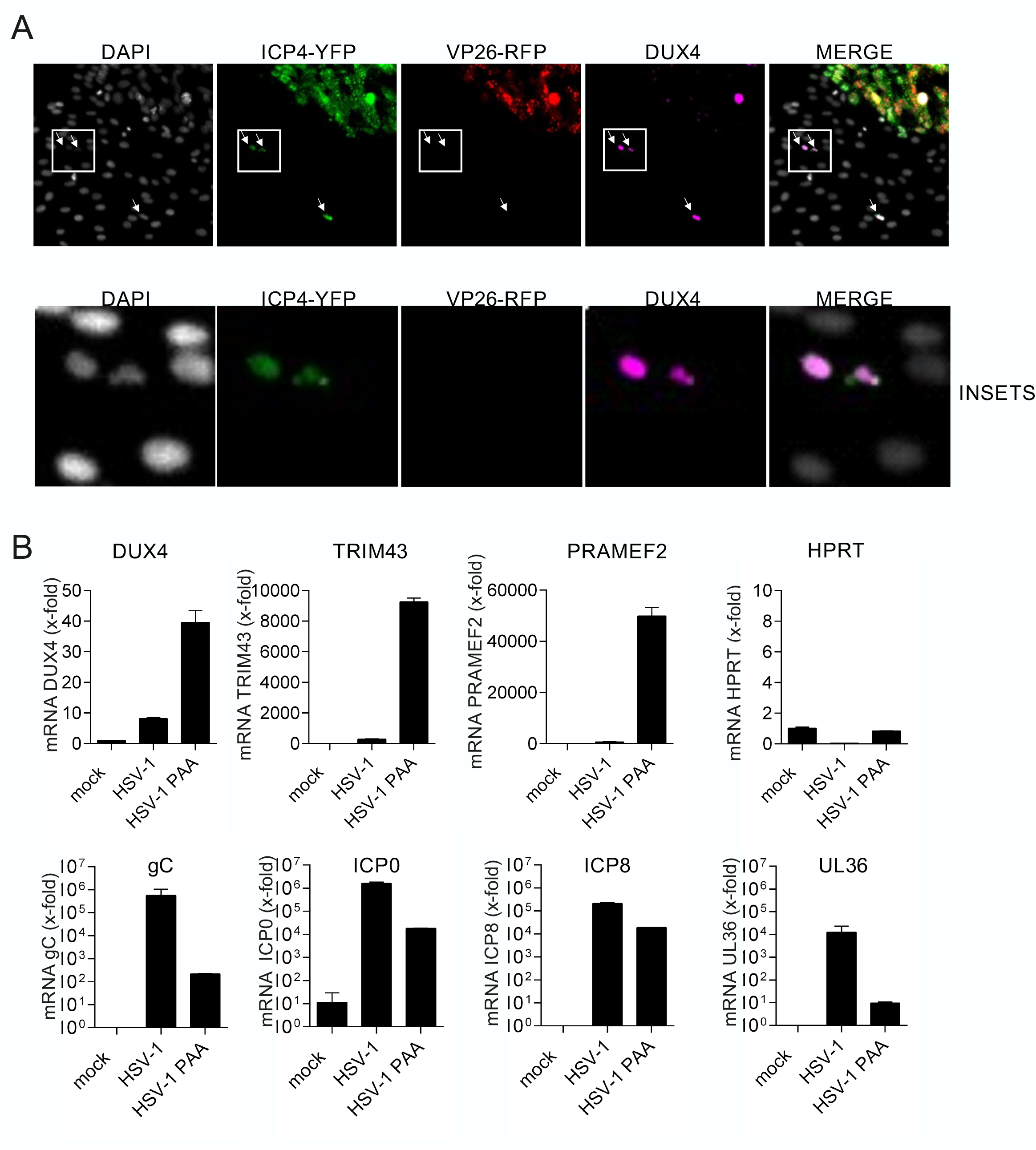
**A**. Immunofluorescence of HDF-TERT cells infected with a recombinant HSV-1 expressing ICP4-YFP and VP26-RFP. DUX4 protein was stained with a DUX4 specific antibody, nuclei were counterstained with DAPI. Nuclear DUX4 expression is only detectable in ICP4+/VP26-cells. B. qRT-PCR analysis of cellular genes DUX4, TRIM43, PRAMEF2 and HPRT as well as viral genes gC, ICP0, ICP8 and UL36 in HDF-TERT cells untreated or treated with PAA and infected with HSV-1 (MOI 0.1). Values are presented as fold induction (normalized to HPRT RNA) relative to uninfected control cells.

It is known that the physiological function of DUX4 during embryonic genome activation at the 2-8 cell stage is to directly bind to DNA and activate genes and retroelements that are necessary for developmental progression^9,15^. In particular endogenous retroelements act as promoters for downstream genes, and binding of DUX4 to retroelements activates transcription of respective genes^18,39^. In order to further analyze the role of DUX4 in herpesviral replication, we performed endogenous DUX4 ChIP-Seq experiments. HFF cells were infected with HSV-1 for 8h and 16h, and binding of DUX4 to both the host and viral genome analyzed. The ChIP-Seq experiments revealed about 11000 DUX4 binding sites within the host genome. Analysis of the binding sites from our endogenous DUX4 ChIP-Seq and comparison with the DUX4 ChIP-Seq performed by Young et al.^18^ with overexpressed DUX4 protein in muscle cells showed almost identical consensus binding sites (Fig. 4A). We observed a strong correlation between DUX4 binding sites at 8h and 14h time points (Fig. 4B). Surprisingly, most DUX4 binding sites were not at transcriptional start sites or within exon regions of genes, but within intronic and intergenic regions (Fig. 4C). Comparison of DUX4-ChIP-Seq binding sites upon HSV-1 infection with RNA-Seq data published from Rutkowski et al. showed that a subset of genes with DUX4 binding sites get upregulated during the course of infection whereas other genes with DUX4 binding sites are either not regulated or even get downregulated upon infection (Fig. 4D). In depth analysis of intronic / intergenic binding sites showed a strong preference of DUX4 binding to repetitive genetic elements (Fig 4E). We found predominant binding of DUX4 to long-terminal repeat (LTR-) elements and short interspersed nuclear elements (SINE) and only a small fraction of binding sites in actual genes (Fig. 4E). During early embryonic development and in particular EGA, it is known that activation of endogenous retroelements can generate alternative promoters for expression of genes, resulting in transcription of developmental genes that are essential for further embryonic development^16,17^. Comparison of DUX4 binding during EGA and herpesviral infection shows that a specific subset of retroelements is bound in both settings (Fig. 4F). Accordingly, reanalysis of RNA-seq data showed a DUX4-mediated activation of a specific subset of LTR-elements that are expressed during EGA. Both HSV-1 infection as well as DUX4 overexpression leads to an induction of the MLT- and THE-class of retroelements, indicating that the expression upon herpesviral infection is driven by DUX4. In addition, HSV-1 infection also induces the LTR12C class of retroelements. Expression of LTR12C retroelements is independent of DUX4 (Fig. 4F), and results in the expression of the previously described C10rf159 antisense transcript^31^, since this transcript starts at an LTR12C element within the first intron of its host gene. Moreover, comparison of genes harboring DUX4 binding sites showed a partial overlap between genes expressed during herpesviral infection and genes expressed during EGA (Fig. 4G)^40^. This indicates that herpesviral induction of the germline-specific transcription factor DUX4 activates transcription of a subset of EGA-specific genes and retroelements. In contrast to the cellular genome, we could not identify robust binding of DUX4 to the HSV-1 genome. The DUX4-ChIP conditions optimized for DUX4 binding to the cellular genome resulted in a high background for the viral genome, most likely due to different physical properties that affect sonication and differences in chromatin accessibility. However, we cannot fully exclude direct binding of DUX4 to the viral genome. We observed a weak potential DUX4 binding site in the promoter region of the viral ICP27 gene, which also contains a potential DUX4 consensus-binding site, and conventional ChIP-experiments followed by quantitative PCR showed an enrichment of DUX4 compared to controls (Fig. S6), suggesting at direct binding of DUX4 to the ICP27 promoter region.

**Figure 4:**
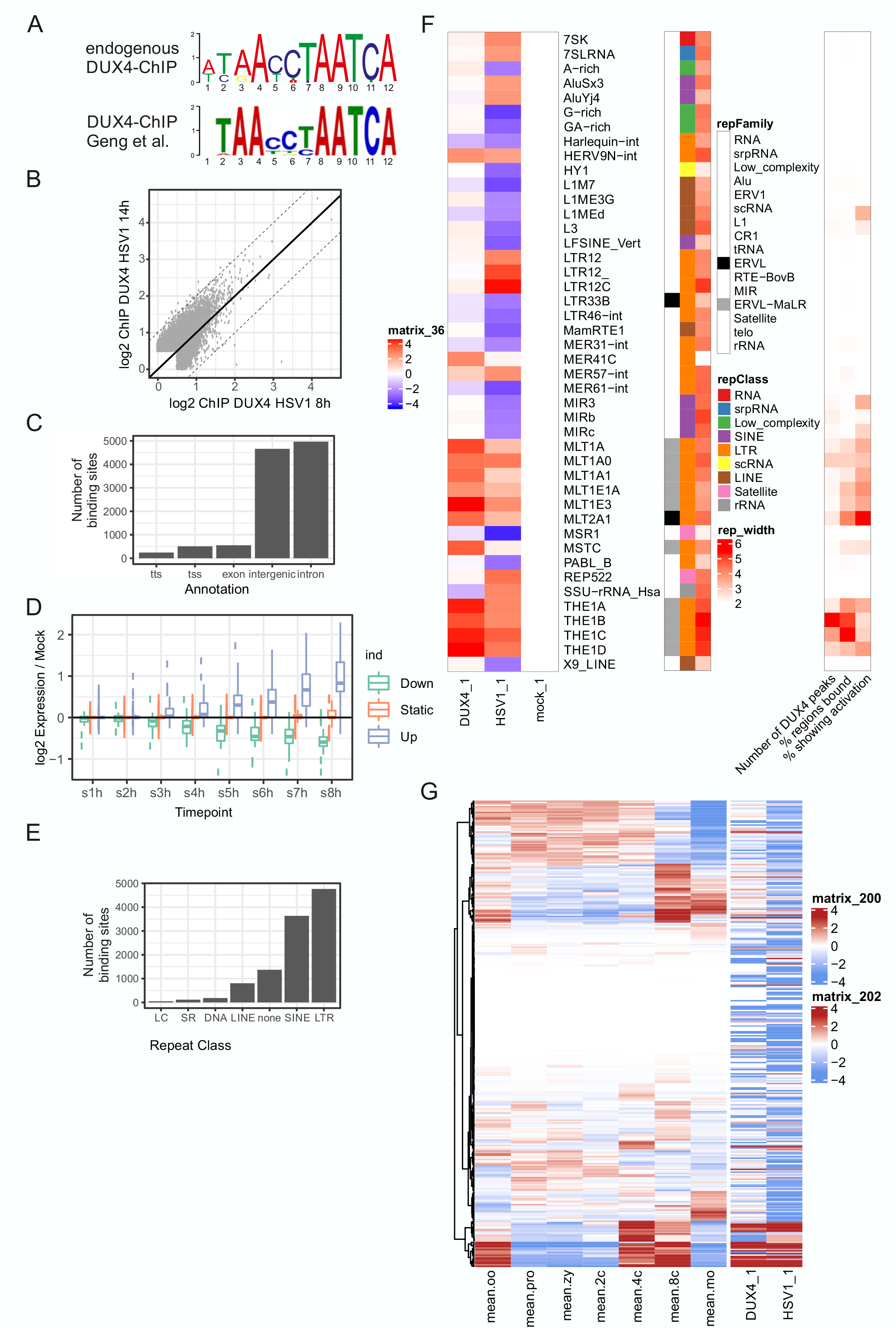
ChIP-Seq analysis of endogenous DUX4 induced by HSV-1 infection **A.-E.** ChIP-Seq of endogenous DUX4 in HFF cells infected with HSV-1 at 8 hpi and 14 hpi. **A.** Comparison of DUX4 consensus binding sequence in HSV-1 infected cells (upper panel) with the previously, published DUX4 consensus sequences (Geng et al.) in the lower panel. **B.** 8 hpi ChIP samples plotted against 14 hpi samples. Both are highly concordant. DUX4 binding increases at 14 hpi. **C.** Number of binding sites in different genomic regions. (TTS: transcription termination site site, TSS: transcription start site) **D.** Different expression levels of DUX4 target genes during HSV-1 infection plotted over time, analysed from Rutkowski et al. 2015 **E.** Analysis of DUX4 binding sites that can be found within repetitive elements. (LC: low complexity repeats, SR: simple repeat) **F.** Repetitive elements regulated in DUX4 overexpressing cells compared to HSV-1 infected cells reanalysed from Full et al. 2019. On the right site repetitive elements are ascribed to their respective class and family. DUX4 binding is scaled to repeat size, number of peaks and activation. **G.** Expression pattern of embryonic genes during embryonic genome activation compared to expression pattern in DUX4 overexpressing and HSV-1 infected cells. Embryonic data was published in (reference) and is normalized to average expression value of each gene. Data from DUX4 overexpression and HSV-1 infection is relative to mock. Shown are genes which have DUX4 binding sites in the proximity of the TSS. (oo: oocyte, pro: pronucelus, zy: zygote, 2c: 2-cell state, 4c: 4-cell state, 8c: 8-cell state, mo: morula)

**Figure 5:**
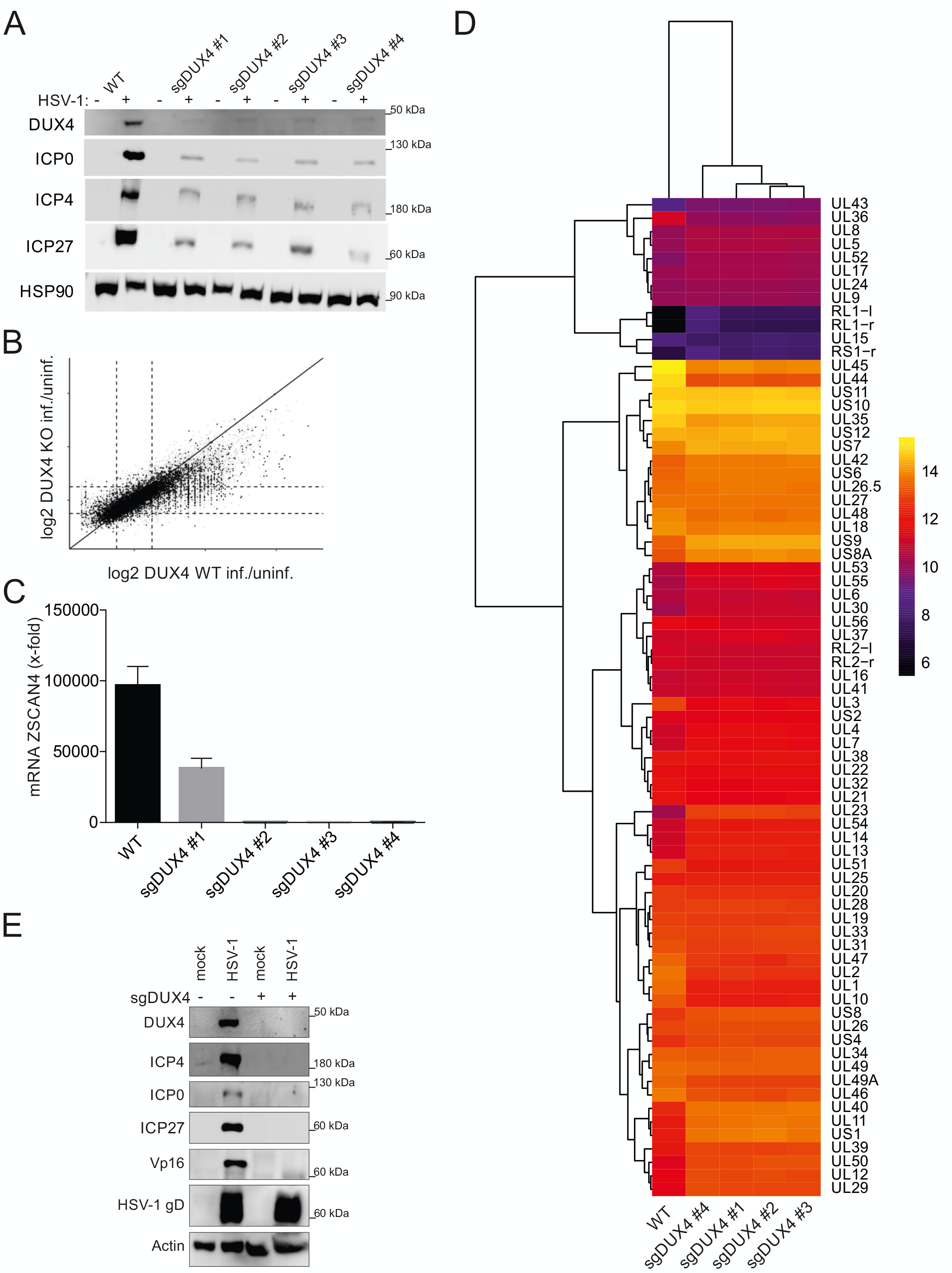
DUX4 expression is critical for herpesviral immediate early protein expression **A.** DUX4 kd cells were generated with sgRNAs #1-4. HSV-1 protein expression in 293T WT cells and 293T cells with DUX4 KD. Cells were infected with HSV-1 (MOI 5) and analysed by western blot. Time points as indicated in graph. Representative experiment out of n=3. **B.** mRNA expression (RNA-Seq.) of DUX4 kd cells plotted against wildtype 293T cells. **C.** qPCR of ZSCAN4 mRNA expression in 293T WT and 293T DUX4 KD cells infected with HSV-1 (MOI 5) for 18h. **D.** RNA-Seq analysis of DUX4 CRISPR/Cas9 treated 293T cells and 293T WT cells infected with HSV-1 (MOI 5) for 18 h. The heat-map illustrates differences in the expression levels of HSV-1 RNAs in WT and KD cells. DUX4 KD cells generated with sgRNAs #1-4. mRNA expression is normalized to expression of vault RNA (VTRNA). One representative experiment out of n=3. **E.** Western blot of 293T WT cells and 293T DUX4 KO cells infected with HSV-1 for 18 h. Comparison of HSV-1 protein expression in WT cells and cells with complete DUX4 KO. Representative experiment out of n=5.

Our data revealed a robust expression of DUX4 during lytic replication of all human herpesvirus subfamilies and thus we wanted to address the physiological consequences in regard to herpesviral gene expression and replication. DUX4 is located within the 4q35 region of chromosome 4^25,26^ and the gene is particularly complicated to target by CRISPR/Cas9. In healthy individuals the locus consists of 10-100 repeat units, and a copy of DUX4 resides in every unit^27^. However, only one of the D4Z4 repeat arrays, the most distal one adjacent to the telomeres is encoding for the functional protein^21^. In addition to multiple repeats on chromosomes 4 (4q allele) there is also another heterochromatic repeat array on chromosome 10 (10q allele)^41,42^ which is likely to interfere with CRISPR/CAS9 based knockout strategies. We therefore used a transient approach in 293T cells to target DUX4 at the population level using 4 different guide RNAs targeting different sequences within the DUX4 locus. The guide RNAs had been successfully used to silence DUX4 in FSHD myocytes previously^48^. We observed a knockdown but not complete knockout of DUX4 upon HSV-1 infection by Western analysis one week after selection with puromycin (Fig. 4A). The knockdown of DUX4 resulted in reduced protein expression of HSV-1 immediate-early genes ICP0, ICP4, and ICP27 upon infection compared to wildtype-cells (Fig.4A). In order to assess the effect of DUX4 knockdown on the transcription of the entire viral genome, RNA-Seq from HSV-1 infected knockdown and wildtype cells was performed. Unsupervised clustering of wildtype and knockdown cell lines showed altered expression of most HSV-1 genes in knockdown cells (Fig. 4B). Whereas the expression of early genes like UL23 and UL54 is lower in wildtype cells, the expression of late genes like UL44/UL45 is higher, indicating that the viral expression cascade proceeds faster in wildtype cells in the presence of DUX4 (Fig. 4B). Analysis of the cellular transcriptome confirmed a reduction of DUX4 target genes in knockdown cells, both by RNA-Seq analysis and qRT-PCR for ZSCAN4 (Fig.4C, D). In addition, we also observed a decrease in expression of endogenous retroelement upon DUX4 knockdown compared to wildtype cells, demonstrating the dependence on DUX4 (Fig.S7). In order to further assess the role of DUX4 in viral gene expression we optimized our CRISPR/CAS9 protocol and could achieve a complete knockout of DUX4 in 293T cells. Infection of DUX4 knockout cells resulted in an almost complete loss of most HSV-1 genes tested in Western Blot, like ICP0, ICP4, ICP27 and VP16. However, the expression levels of HSV-1 gD were not affected by knockout of DUX4, which is in accordance with published data showing that the expression of gD is independent of the viral expression cascade^49^.

## Discussion

In humans the female oocyte is produced during female gametogenesis in the embryo and is then stored in prophase I of the meiosis for up to 50 years. The oocyte transcription is halted by epigenetic mechanisms, and stored mRNAs mostly control development. After fertilization the zygote is formed, maternal to zygotic transition (MZT) takes place and with the onset of EGA the zygotic genome starts to control transcription^13^. In order to induce transcription, the embryo has to overcome silencing of the genome, which is regulated by repressive epigenetic modifications like DNA methylation, histone modifications and a shortage of the cellular transcription machinery^14,43^. DUX4 has been shown to be essential for EGA by activating hundreds of genes that are necessary for further development, and several endogenous retroelements^9,10,15^. Although the detailed function of most of DUX4 target genes remains elusive, it is thought that DUX4 induces several important factors that are involved in the creation of a permissive environment that allows transcription from the newly formed diploid genome.

Upon herpesviral infection the viral genome enters the nucleus and gets chromatinized, although the degree of chromatinization of the viral genome is discussed controversially, in particular for HSV-1. However, it is widely accepted in the field that herpesviruses are subjected to epigenetic silencing, and that they evolved strategies to prevent epigenetic silencing of their genome in order to allow transcription necessary for viral replication and virus transmission. We show that herpesviruses from all human subfamilies induce a robust expression of DUX4 upon lytic infection. Considering that alpha-, beta-, and gammaherpesviruses were split into three separate lineages about 200 million years ago^44^, this hints at a highly conserved mechanism in the coevolution of herpesviruses with their respective hosts. For HSV-1, we demonstrate that DUX4 expression is induced by viral tegument/immediate-early proteins ICP0 and ICP4, indicating that this is an active induction by herpesviruses. Viral ICP0 and ICP4 induce expression of DUX4 at early stages of infection for a brief period, indicated by nuclear staining of DUX4 in ICP4+/VP26-cells. In ICP4+/VP26+ double positive cells we either observed no DUX4 staining or cytoplasmic DUX4 staining. Blocking viral DNA replication with PAA results in higher levels of DUX4 and its target genes, suggesting DUX4 behaves similarly to the strong HSV-1 beta genes, which are upregulated early in infection and are shut down during the switch to viral DNA replication and late gene expression. Interestingly, this very short induction of DUX4 resembles the mechanism of DUX4 induction during EGA, where a short burst of DUX4 is sufficient for target gene activation and reprogramming of the embryo but prevents toxicity mediated by DUX4.

Moreover, we could demonstrate that herpesviral DUX4 induction is essential for efficient herpesviral gene expression. Depletion of DUX4 from cells results in a loss of most herpesviral protein expression, as shown for HSV-1. We hypothesize that herpesviruses evolved to partially mimic EGA by actively inducing DUX4 expression. EGA is conserved in all animals, and due to its significance in embryonic development there is very little room for the host to antagonize this viral mimicry of EGA^13^. Any interference with DUX4 function, for example by mutations in the coding sequence or by preventing DUX4 expression could lead to drastic changes in the embryonic development that are incompatible with life. Thus, from a viral perspective, it is beneficial to exploit a host gene which is essential for development for its own purpose in order to limit mutations that affect viral replication.

In addition to its role in EGA and in the development of FSHD, it was recently published that DUX4 also plays an important role in a variety of human cancers^45^. Reanalysis of almost 10000 cancer transcriptomes from The Cancer Genome Atlas (TCGA) showed DUX4 reexpression in many human cancers^45^. The authors speculate that expression of DUX4 and DUX4 target genes may contribute to tumorigenesis^45^. Interestingly, the human gammaherpesviruses, Epstein-Barr Virus (EBV) and KSHV are classified as human carcinogens by the WHO and cause cancer in humans^6^. It would be very informative to see but beyond the scope of this manuscript whether DUX4 expression induced by EBV and KSHV also contributes to herpesviral oncogenesis. In addition the authors showed that DUX4 expression results in a downregulation of MHC-Class I expression, hinting at a possible immune evasion mechanism^45^. Although herpesviruses of all subfamilies already encode for several immune evasion genes that interfere with presentation of viral peptides by MHC-I, the downregulation of MHC-I by DUX4 could also contribute to herpesviral immune-evasion that prevents T-cell killing of infected cells *in vivo.*

We could show that herpesviral DUX4 expression also leads to expression of several endogenous retroelements, including LTR-containing retrotransposons, LINE-1 elements and Alu-elements. Whereas cell type specific differences in the induction of retroelements exist, with more retroelements being transcribed in tumor cell lines than in primary cells, we identified an Alu-element 5’ of the ZSCAN4 gene that serves as a binding site for DUX4 and drives ZSCAN4 expression in all cell lines tested. In addition it is known for years that herpesviral infection leads to the induction of endogenous HERV-W retroviruses and also to the reactivation of the HIV-LTR^46,47^, hinting at a role of herpesviral DUX4 in respective processes. It is hypothesized for years that some herpesviruses including HSV-1 have evolved a high GC-content in their genome in order to prevent insertion of endogenous retroelements that have a bias for AT-rich sequences. The active induction of DUX4 by HSV-1 proteins and the importance of DUX4 for herpesviral gene expression / replication supports this possible explanation for the high GC-content of HSV-1 genomes. It may help to preserve herpesviral genome integrity despite the DUX4-mediated induction of endogenous retroelements that could integrate into the viral genome.

Our data points at a very important if not essential role for DUX4 in herpesviral gene expression and replication. We could demonstrate that depletion of DUX4 in cells almost completely abrogates viral immediate early gene expression. As such it is tempting to speculate that DUX4 and DUX4 downstream genes could be targeted for therapy of herpesvirus-associated diseases. Since aberrant DUX4 expression is the cause of FSHD, several drugs targeting DUX4 are currently under development using a variety of different strategies. Blocking DUX4 means preventing herpesviral immediate early gene expression and subsequent viral replication from the beginning. Targeting a cellular protein has the major advantage that viral escape mutants and resistance-formation are far more unlikely. Therefore, DUX4 provides an attractive target for anti-herpesviral therapy.

## Supporting information

Walter et al Supplemental Figures S1-S7

## Acknowledgments

We are grateful to David M. Knipe (Harvard Medical School, Boston, USA), Benedikt Kauffer (Freie Universität Berlin, Berlin, Germany), Alexander Steinkasserer (University Hospital Erlangen, Germany), Roger Everett (University of Glasgow, Scotland), Adam Whisnant and Lars Dölken (University of Würzburg, Germany) for providing plasmids and virus stocks. We also thank Dr. Benjamin Schmid and Dr. Philipp Tripal the Optical Imaging Center Erlangen (OICE) from the Friedrich-Alexander University Erlangen for support with microscopy. This study was supported by grants from the German Research Foundation (DFG En 423/5-1, to AE), the IZKF Erlangen (J57, to FF), the BMBF Junior Research Group “Duxdrugs” (01KI2017, to FF) and the NIH grant GM127527 (S.T.).

## Methods

### Cell culture and viruses

HEK 293T, primary HFF, Vero were maintained in Dulbecco’s modified Eagle’s medium (DMEM, Thermo Fisher Scientific) supplemented with 10% (vol/vol) heat-inactivated fetal bovine serum (FBS, Thermo Fisher Scientific), 2 mM GlutaMAX (Thermo Fisher Scientific), 1 mM HEPES (Thermo Fisher Scientific) and 1% gentamycin (vol/vol). iSLK harbouring recombinant (r) KSHV.219 were cultured in DMEM supplemented with 10% (vol/vol) FBS, 2 mM GlutaMAX (Thermo Fisher Scientific), 10 mM HEPES buffer (Thermo Fisher Scientific) and 1% penicillin-streptomycin (vol/vol), 1 μg/ml Puromycin and 250 μg/ml C418. All cells were kept under standard culture conditions and regularly checked for mycoplasma contamination (MycoAlert Kit, Lonza).

Lytic replication of KSHV in iSLK.219 was induced by adding 1 μg/ml doxycycline. Infection experiments were performed with HSV-1 strain KOS, HSV-1 GFP (F-strain, provided by B. Kauffer, Berlin) or HCMV (strain Ad-169). HSV-1 deltaICP0 (ICP4-YFP) and HSV-1 delta ICP27 were provided from R. Everett (University of Glasgow). HSV-1 T-VEC del ICP34.5/ICP47 IMLYGIC^®^, Talimogene laherparepvec) was provided by L. Heinzerling (Friedrich-Alexander-Universität Erlangen-Nürnberg). HSV-1 expressing ICP4-YFP and VP26-RFP is a new recombinant virus generated in the background of strain 17 by recombining HSV-1 expressing ICP4-YFP (obtained from Matthew D. Weitzman) and HSV-1 expressing VP26-RFP (obtained from Oren Kobiler) and plaque purifying for 5 cycles.

### Reagents, plasmids and transfections

Transfections were performed with GenJet (SignaGen Laboratories) or Lipofectamin 2000 (Thermo Fisher Scientific) according to the manufacturer’s protocol. Plasmids are listed in Table 3. KU60019 was purchased from Selleckchem.

**Table 1.**
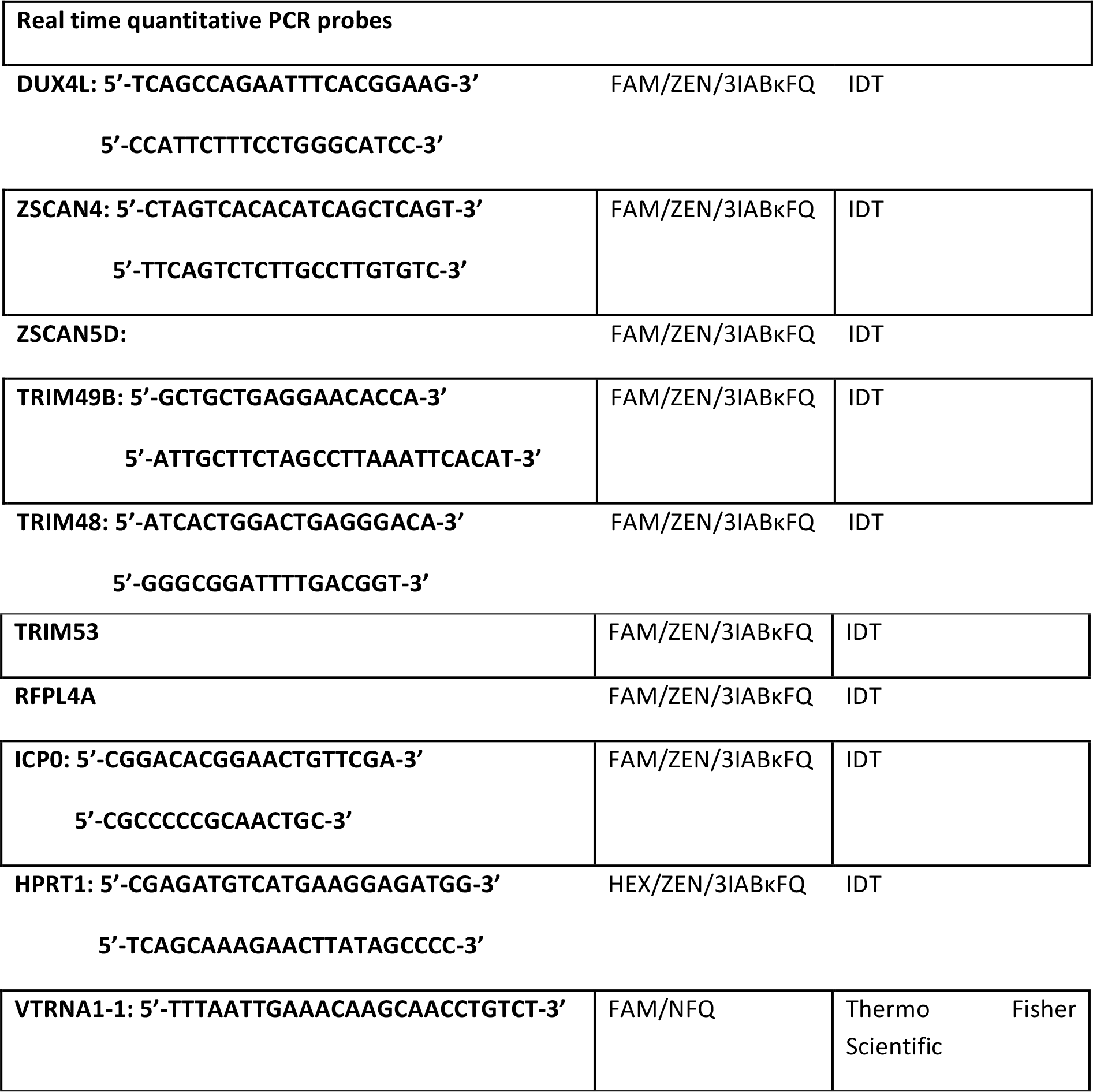
Oligonucleotides

**Table 2.**
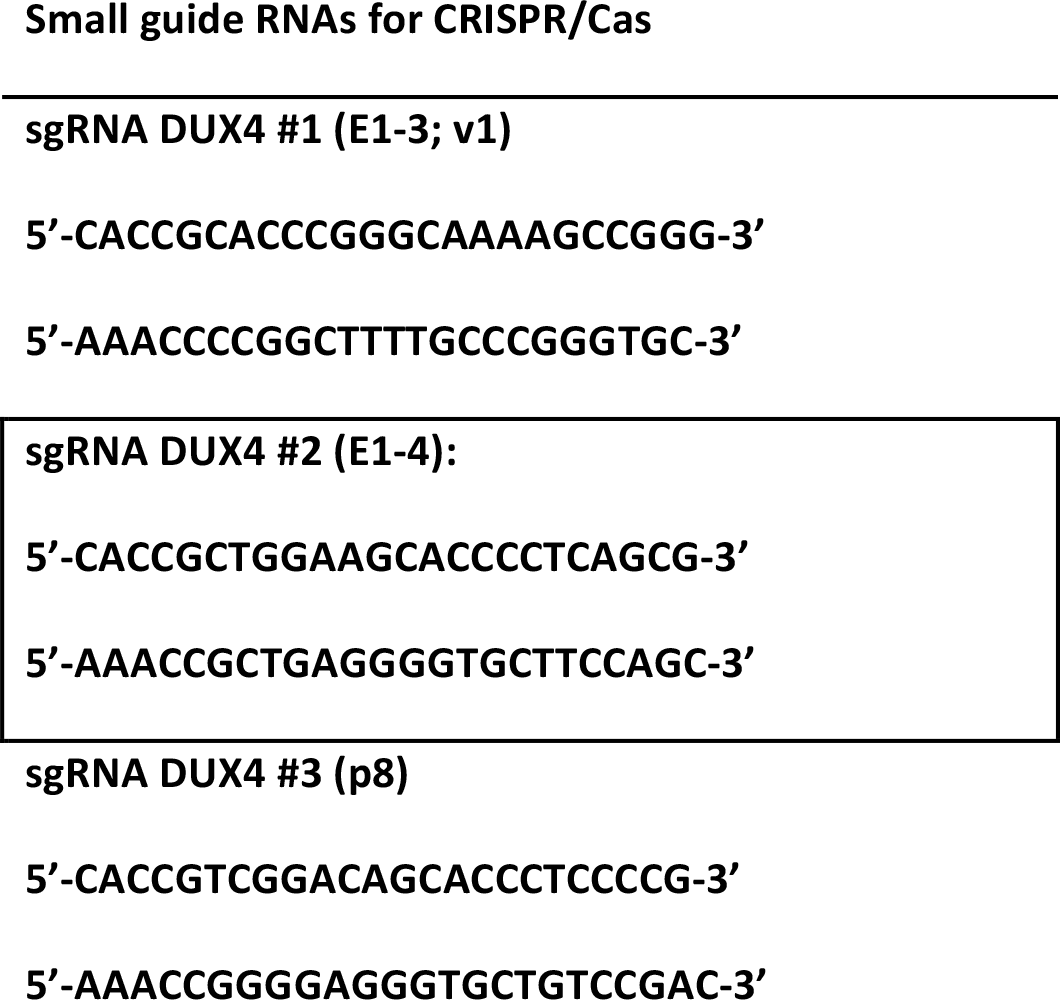
gRNAs

**Table 3.**
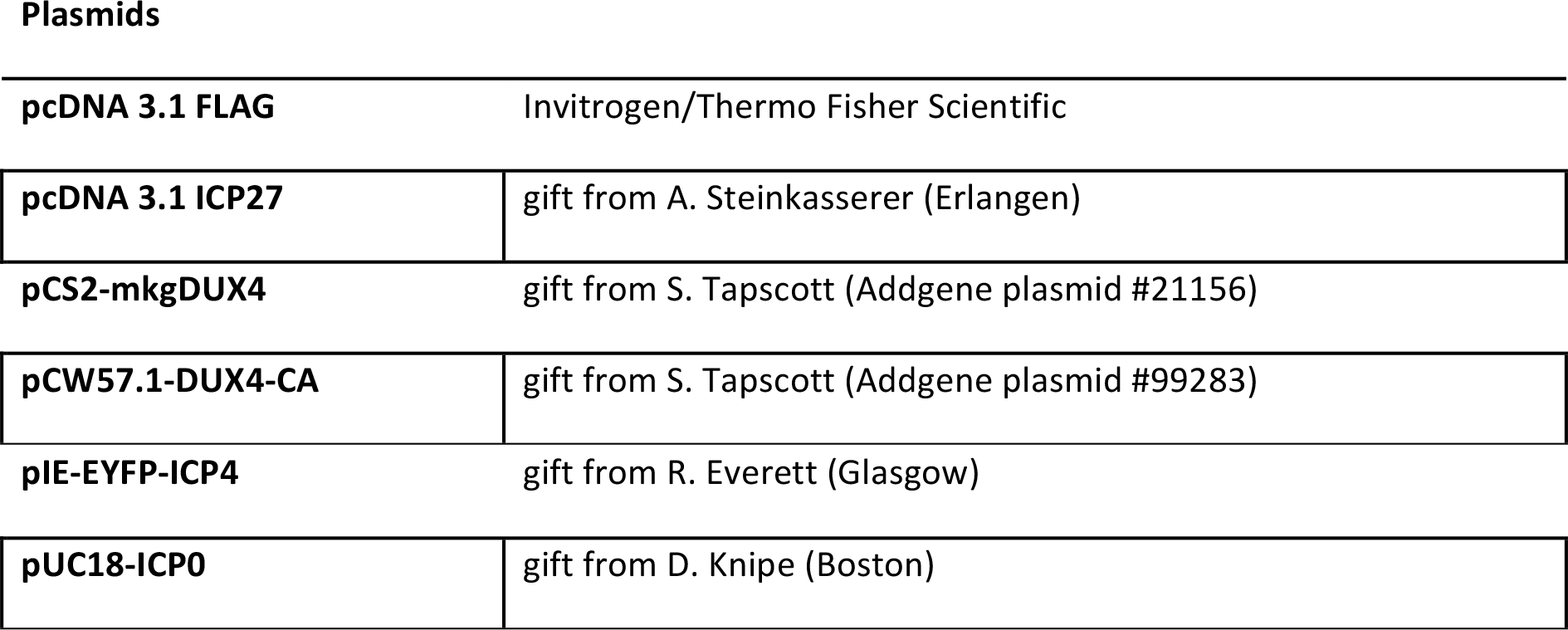
Plasmids

### Western Blotting

Cells were lysed in RIPA HS buffer (10 mM Tris-HCl pH 8.0, 1 mM EDTA, 500 mM NaCl, 1% Triton X-100 (vol/vol), 0,1% SDS (vol/vol), 0,1% deoxycholic acid (DOC) with Aprotinin and Leupeptin, MG-132 and sodium metavanadate (Sigma-Aldrich). The cell pellet was centrifuged at 4°C, 14.000 rpm for 30 min. Samples were diluted with Laemmli-SDS sample buffer and heated for 5 min at 95°C. Antibodies used are listed in Table 4.

**Table 4.**
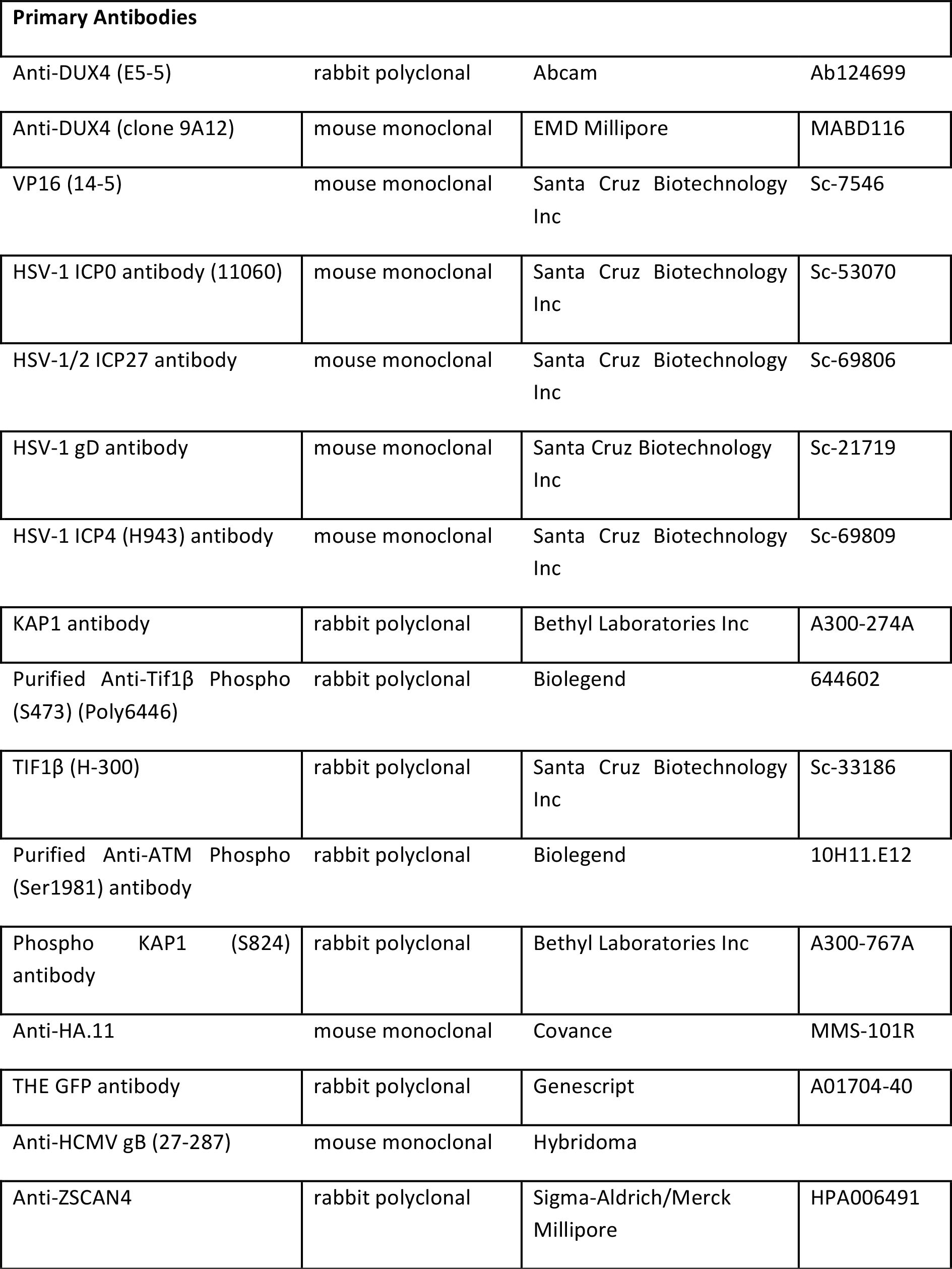
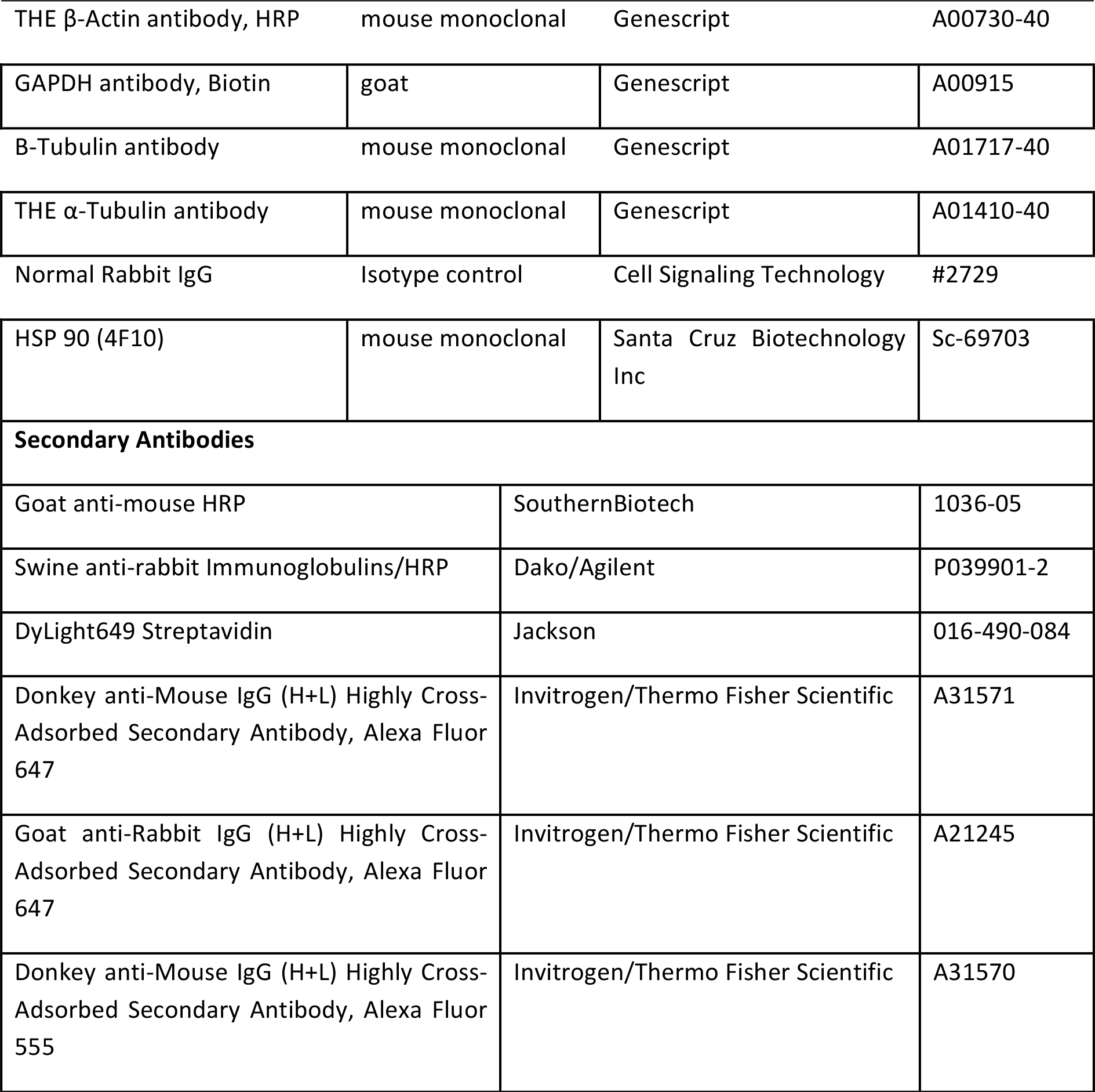
Antibodies

### qRT-PCR

RNA was extracted using the Direct-zol RNA Miniprep Plus kit from Zymo Research according to manufacturer’s instructions. Reverse transcription was performed using the Super Script IV Kit (Thermo Fisher Scientific) according to manufacturer’s instructions. qRT-PCR was carried out using TaqMan^™^ Universal PCR Master Mix I (Applied Biosystems, Thermo Fisher Scientific) 0,1 μg as template on a 7500 Fast Real-Time PCR machine. Or reverse transcription and qRT-PCR were conducted in one step using 0.1 μg RNA as template with the Luna Universal Probe One-Step RT-qPCR kit (New England Biolabs) following manufacturer’s protocols. Primers/probes are listed in Table 1. Expression levels for each gene were obtained by normalizing values to *HPRT1* or *VTRNA* and fold induction was calculated using the comparative CT method (ΔΔCT method).

### CRISPR and sgRNAs

All sgRNAs used in this study were previously described^48^ and are listed in Table 2. sgRNAs were cloned into LentiCRISPRv2 plasmid gifted from F. Zhang (Addgene plasmid #52961) and verified by sequencing. Lentiviruses were packaged with pMD2.G (Addgene plasmid #12259) and psPAX2 (Addgene plasmid #12260) (both gifted from D. Trono) into HEK 293T. HEK 293T were seeded into 12 well plates and lentiviral supernatants added at 70-80% confluency. Plates were centrifuged at 1200 rpm for 2 min after centrifugation culture medium was added and cells incubated over night at 37°C. The medium was changed to normal culture medium the next day and selection with 2 μg/ml puromycin in normal culture medium started on day 3.

### Next generation Chip-Seq

For CHIPmentation primary HFF cells were seeded in T175 flasks and infected with HSV-1 KOS (MOI 10) for different time points. CHIPmentation was conducted as previously described (Schmidl et al. 2015). Cells were sonicated using the Bioruptor (Diagenode) for 30 cycles. For the immunoprecipitation protein G Dynabeads (Thermo Fisher Scientific) were used. Samples were incubated with either 2.5 μg Anti-DUX4 (E5-5) (Abcam) or Normal Rabbit IgG (Cell Signaling Technology) as control. Samples were purified using AMpureXP beads (Beckman Coulter) according to manufacturer’s description.

### Next generation RNA-Seq

HEK 293T wildtype cells and HEK 293T CRISPR/Cas knockout cells were seeded in T25 flasks. Cells infected for different time points with HSV-1 KOS (MOI 10). Cells were lysed in Trizol (Life Technologies by Thermo Fisher Scientific), and total RNA was isolated using the RNA clean and concentrator kit (Zymo Research), according to the manufacturer’s instructions. Sequencing libraries were prepared using the NEBNext Ultra II Directional RNA Library Prep Kit for Illumina (NEB) with 9 cycles PCR amplification, and sequenced on a HiSeq 4000 device with 1×50 cycles. For quantification of viral gene expression alignments were done using hisat2^49^ on the HSV-1 genome (strain 17, genbank accession no. NC_001806) and readcounts per gene quantified using quasR^51^.

### Bioinformatic analysis of ChlP-seq data

ChIP-seq data processing was done using the PiGx-ChIP-seq pipeline (https://doi.org/10.1093/gigascience/giy123). In short, adapters and low quality bases were trimmed from reads using Trim-galore. The reads were mapped on the hg19 version of the human genome, combined with HSV-1 genome, using Bowtie2 (22388286) with k = 1 parameter. bigWig tracks were created by extending reads to 200, collapsing them into pileups, and normalizing to reads per million. Peak calling was done with MACS2 (https://github.com/taoliu/MACS) using the default parameters. Motif discovery was done using MEME (25953851) with the default parameters, on the top 100 peaks (sorted by q value), in a region of +/- 50bp around the peak center. Peak annotation was done using the hg 19 ENSEMBL GTF file, downloaded on 17.03.2017. from the ENSEMBL database (29155950). Peaks were annotated based on the following hierarchy of functional categories: tss -> exon -> intron -> intergenic (eg. if a peak overlapped multiple categories, it was annotated by the class that is highest in the hierarchy). Peaks were overlapped with the hg19 Repeatmasker repeat annotation, downloaded from the UCSC database (30407534) on 03.02.2015.

### Transposon Quantification for KD samples

Transposon expression was quantified using TeTranscripts (26206304) version 2.2.0, using the hg19 version of the human genome and a custom formatted transposon GTF file, made available by the Hammell laboratory from the following link: http://labshare.cshl.edu/shares/mhammelllab/www-data/TEtranscripts/TE_GTF/. Before visualization, the read counts per TF repName repeatmasker category were multiplied by the following size factors 10000/(number of repeats), and 1000/(average repeat length). The transposon expression was visualized relative to the uninfected wild type sample, using the ComplexHeatmap package (27207943).

### Comparison of HSV-1 gene expression with the embryonic expression profile

Expression profiles for HSV-1 and DUX4 ectopically expressed, Hela cells were taken from the following publication (30420784). The embryonic expression profiles were downloaded from the ARCHS4 database (29636450). The data originated from the following repository GSE44183. Data was visualized using the ComplexHeatmap function.

### Comparison of repeat expression during HSV-1 infection and DUX4 overexpression

Expression profiles obtained from HSV-1 and DUX4 ectopically expressed, Hela cells were taken from the following publication (30420784). DUX4 binding data extracted from the supplementary data from the following publication 24278031. RNA - seq data was mapped using STAR, and the repeats were quantified by counting the number of, uniquely mapping, spliced reads overlapping with the each transposon category. Expression was visualized using ComplexHeatmap. Prior to the visualization the reads were normalized to uninfected samples.

### Statistical analysis

*P* values were calculated using an unpaired Student’s test. *P* <0.05 was considered statistically significant.

