## Supplementary material for "Herpesviral induction of germline transcription factor DUX4 is critical for viral gene expression": Walter et al Supplemental Figures S1-S7

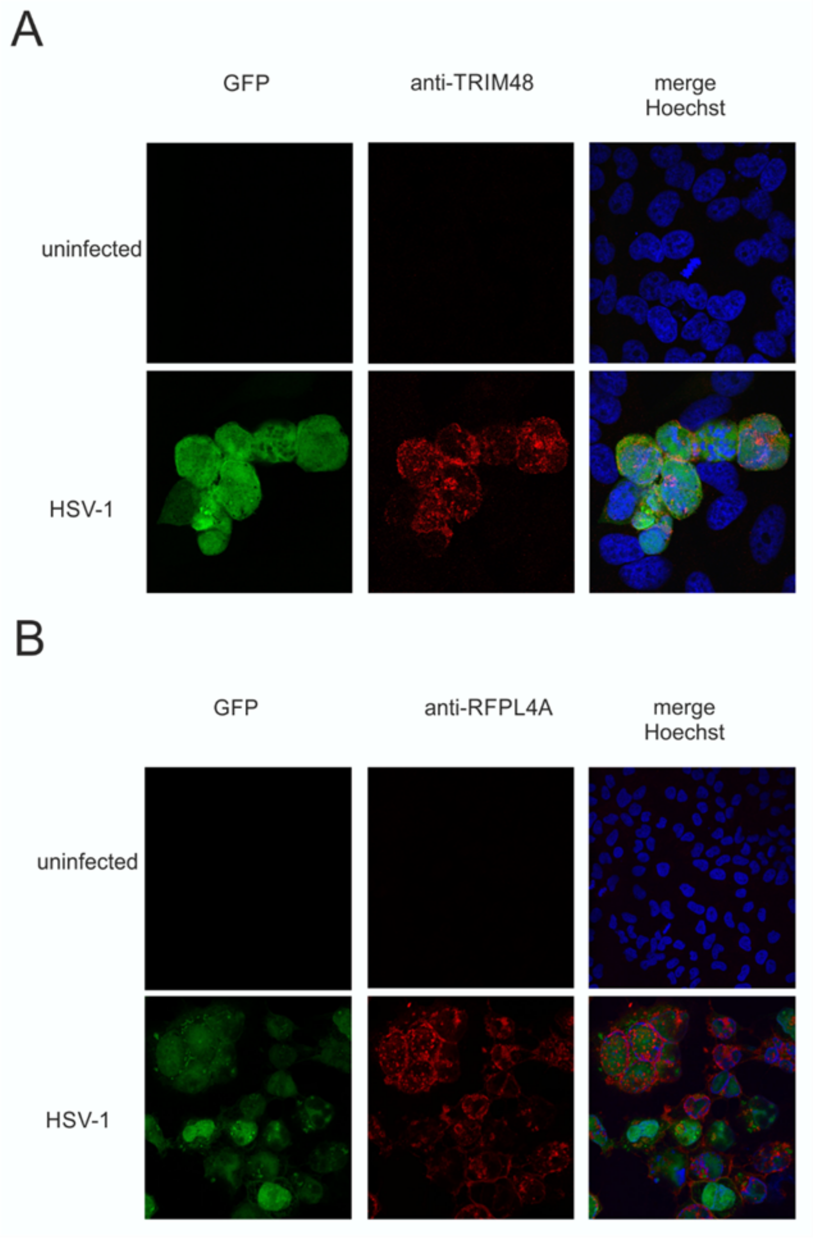


**Fig. S1: Expression of RFPL4A and TRIM48 upon HSV-1 infection.**
Immunofluorescence of HeLa cells infected with green fluorescent protein- (GFP-) tagged HSV-1 for 18 h. One representative out of n=3. Nuclei are stained with Hoechst 33342 in blue. **A.** Cells were stained with anti-TRIM48 antibody in red. **B.** Cells were stained with anti-RFPL4A antibody in red.

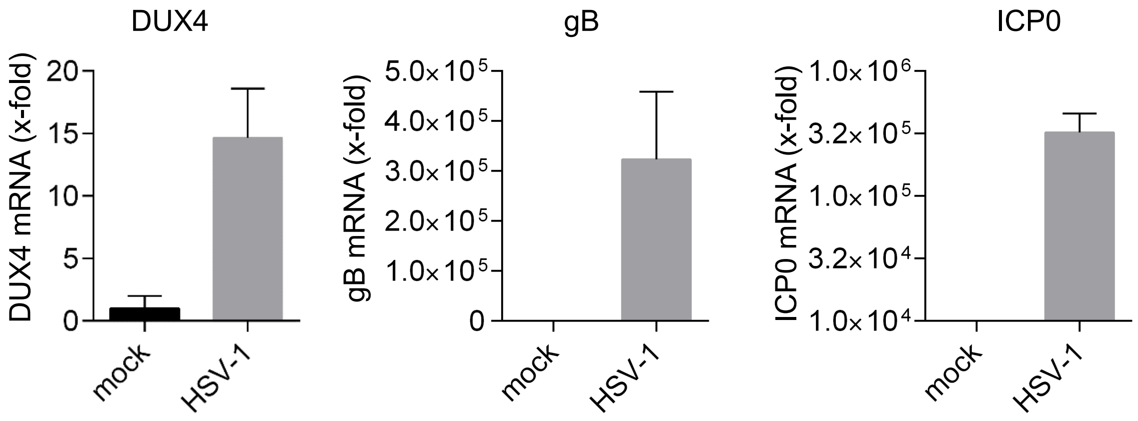


**Fig. S2. DUX4 expression is upregulated in primary human melanocytes** qRT-PCR of cells infected with HSV-1 for 18 h (MOI 0.1). HSV-1 ICP0 and gB served as infection controls. Data represent mean and s.d. of *n* = 3 (biological replicates)

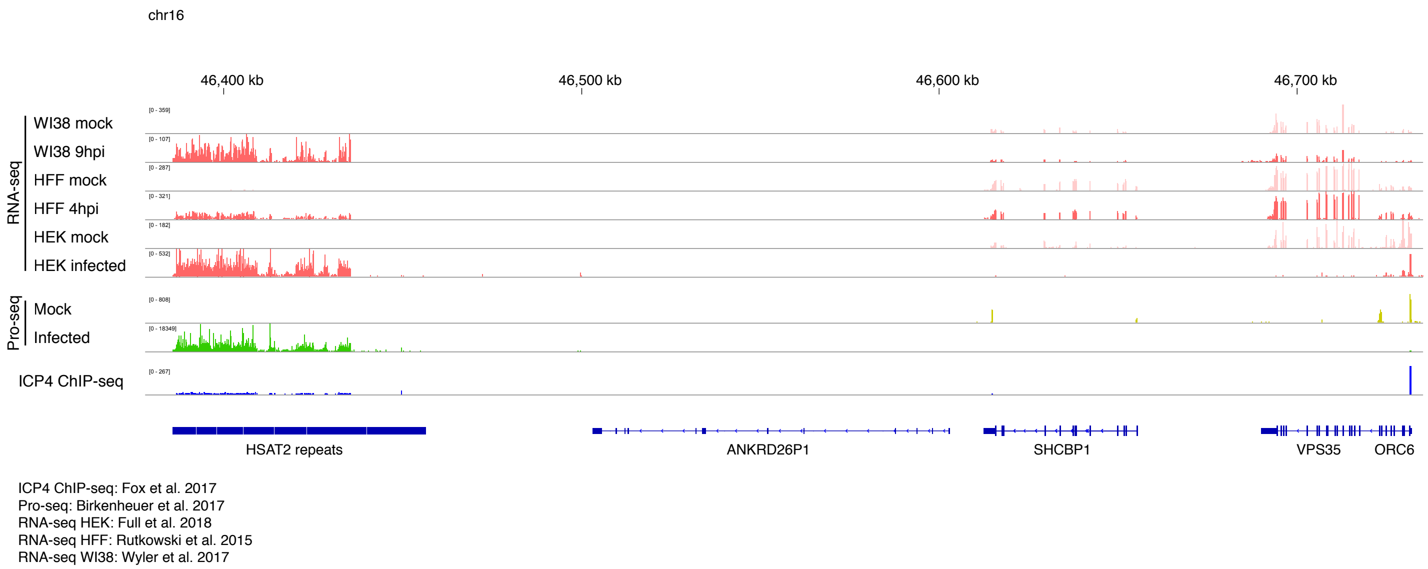


**Fig. S3. Upregulation of HSATII repeats upon HSV-1 infection** RNA-Seq. data of the HSATII repeats and neighbouring loci in WI38 9 hpi, HFF 4 hpi and HEK 293T cells 18 hpi. Pro-Seq. (Precision Run-On Sequencing ) shows RNA-pol II occupancy at HSATII repeats.

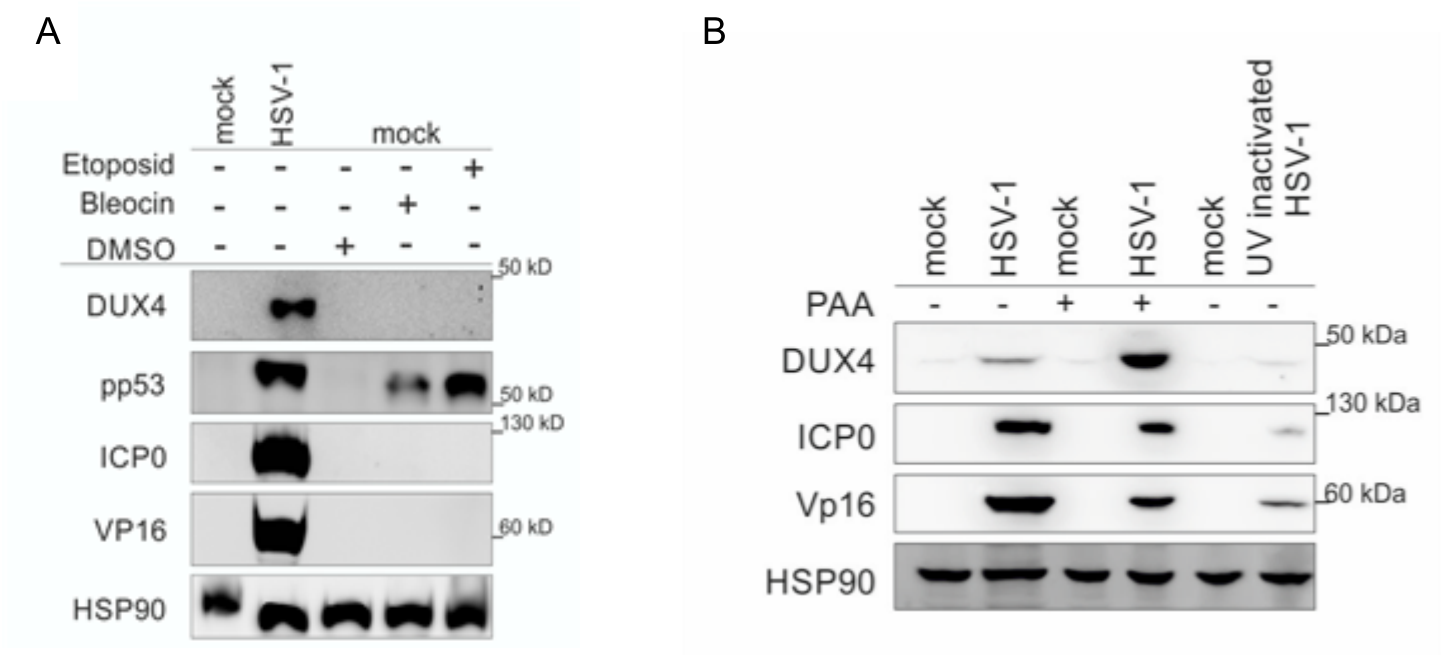


**Fig. S4. DUX4 expression is independent of the DNA damage response (DDR) and viral DNA replication. A** Western blot of DUX4 expression in 239T cells infected with HSV-1 for 18 h or treated with Bleocin or Etoposid. Bleocin and Etoposid were used to induce DNA damage in the cells and start the DDR. HSV-1 ICP0 and Vp16 were used as markers for infection. HSP90 was used as loading control. One representative out of n=3 **B**

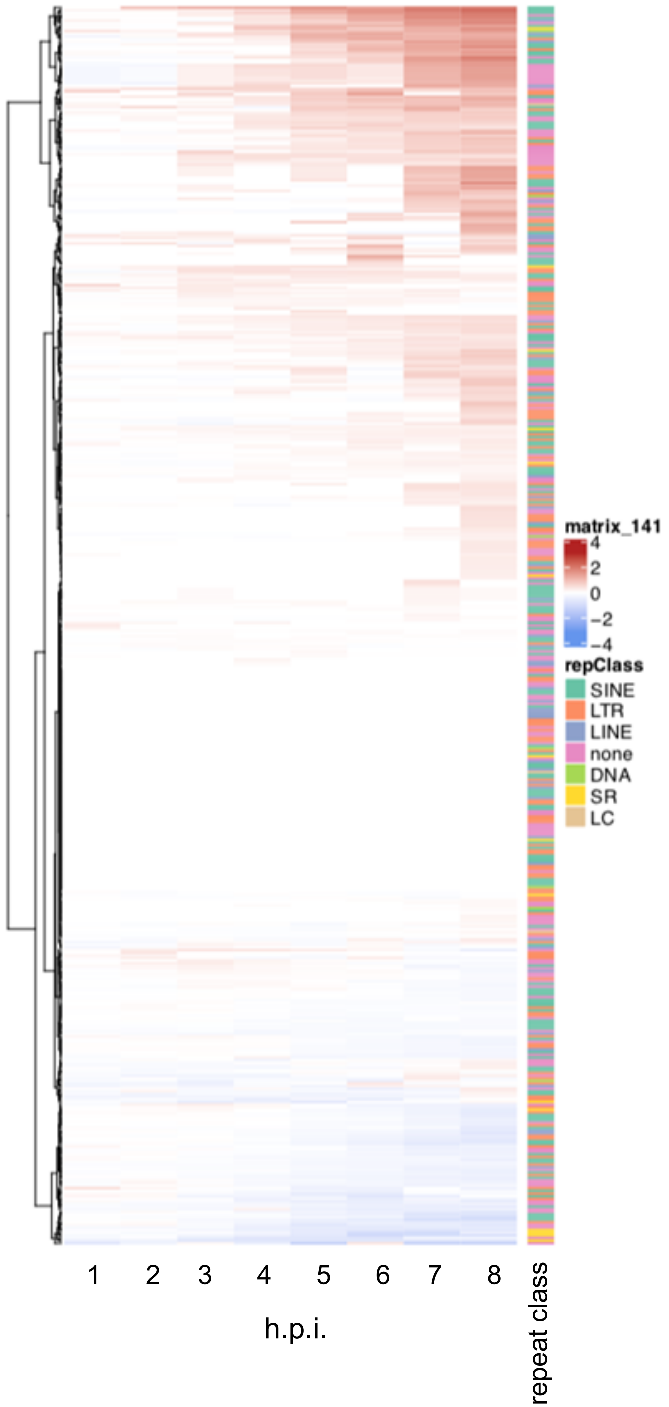


**Fig. S5. DUX4 target genes have variable dynamics during infection.** Genes are upregulated / downregulated with different kinetics (based on 4sU data by Rutkowski et al.). Whereas some genes are upregulated quite early, the bulk of upregulation happens at 3-4 hpi. Upregulated genes are enriched for methylation mediated chromatin silencing proteins, while the downregulated genes are enriched for antigen presentation. The most enriched tissue in the human ARCH4 expression database is the human zygote. The second most expressed gene is OOEP which is an early embryonic protein necessary for progression through the zygotic activation. h.p.i. (hours post infection)

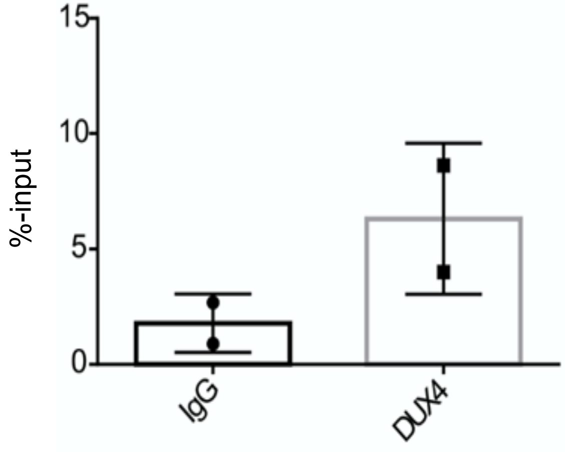


**Fig. S6. Binding of DUX4 to the HSV-1 ICP27 promoter**
ChIP of DUX4 in cells infected with HSV-1 for 8 h MOI 5, analyzed by qPCR for ICP27 promotor binding. Normal IgG was used as antibody control. One representative out of n=2

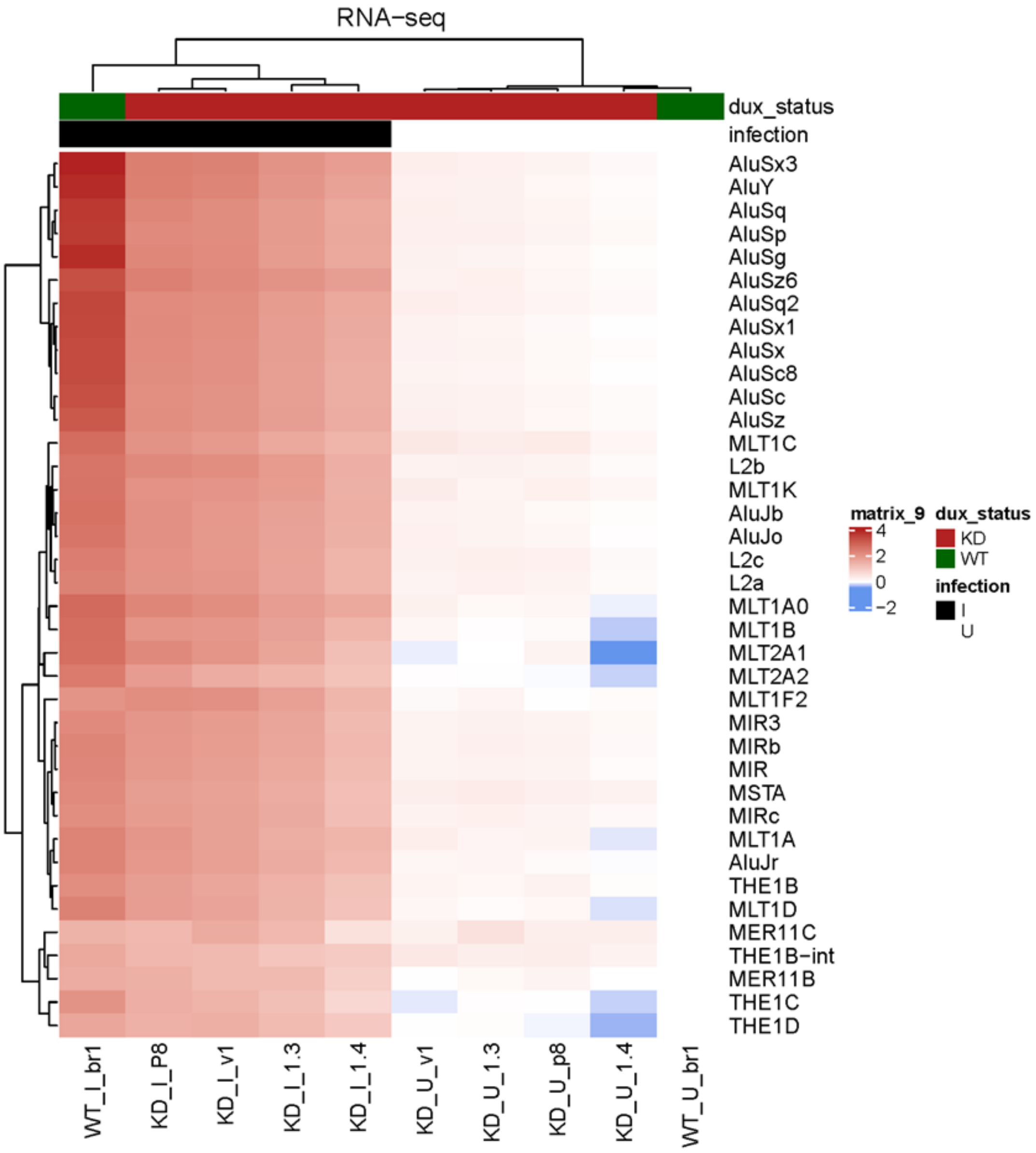


**Fig. S7. Upregulation of endogenous retroelements upon HSV-1 infection is dependent on DUX4.** RNA-Seq of wildtype (WT) and DUX4 knockdown (KD) 293T cells infected with HSV-1. The heatmap shows the expression of endogenous retroelements in cells uninfected (U) or infected (I) with HSV-1.
